# Oral epithelial expression of angiotensin converting enzyme-2: Implications for COVID-19 diagnosis and prognosis

**DOI:** 10.1101/2020.06.22.165035

**Authors:** Mythily Srinivasan, Susan L Zunt, Lawrence I Goldblatt

## Abstract

The severe acute respiratory syndrome coronavirus-2 (SARS-CoV-2) uses the angiotensin converting enzyme (ACE)-2 as the host receptor for target cell entry. The extent and distribution of ACE-2 has been associated with the clinical symptoms of coronavirus disease (COVID)-19. Here we show by immunofluorescence analysis that the ACE2 is abundantly expressed in oral mucosa, particularly in the surface epithelial cells suggesting that these cells could represent sites of entry for SARS-CoV-2. Further, together with the reports on ACE2 ectodomain shedding, we discuss the rationale for the hypothesis that the ACE-2 measurement in saliva could be a marker for COVID-19 infection during early phase following SARS-CoV-2 exposure.

## Introduction

Coronaviruses (CoV) are a group of enveloped viruses with non-segmented, single-stranded, and positive sense RNA genomes. While most coronaviruses cause mild respiratory illnesses, the first epidemic was caused by the severe acute respiratory syndrome (SARS)-CoV-1 in 2003. SARS-CoV-1 spread to over two dozen countries affecting more than 8000 individuals with nearly 10% mortality^1,2^. The current pandemic caused by SARS CoV-2 has spread much more rapidly affecting nearly 7 million individuals globally with over 400,000 deaths as of June 2020.

Both SARS-CoVs encode at least four major structural proteins including the spike protein, the membrane protein, the envelope protein, and the nucleocapsid protein. The spike protein that protrudes from the surface of the virus is a type I glycoprotein that binds specific host cell receptor via a receptor binding domain facilitating viral entry into target cells^3^. The SARS-CoV-1 has been shown to use a metallopeptidase named angiotensin-converting enzyme 2 (ACE2) as a receptor for target cell entry^4,5^. Positive correlation between ACE2 expression and SARS-CoV-1 infection has been observed^6^. Furthermore, ACE2 expression on non-permissive cells conferred susceptibility to SARS-CoV-1 infection, indicating that ACE2 is sufficient to allow viral entry^7,8^. Transgenic mice overexpressing human ACE2 in alveolar epithelial cells and other epithelia developed rapidly progressing respiratory illness following intranasal exposure of SARS-CoV-1^9^.

Genomic and structural studies have shown that the receptor binding domain of SARS-CoV-2 exhibits similar molecular characteristics as that of SARS-CoV-1^10,11^. In vitro cellular studies showed that that the SARS-CoV-2 used ACE2 and no other coronavirus receptors such as aminopeptidase for cellular entry^11^. Indeed SARS-CoV-2 has been shown to bind ACE2 with higher affinity than the SARS-CoV-1^12^. Upon binding ACE2, the spike protein of SARS-CoV-2 is cleaved by host proteases such as the TMPRSS2, thereby releasing the spike fusion peptide and facilitating host cell entry^10,13^. Replication within the target cells establishes the SARS-CoV-2 infection mediating tissue damage.

ACE2 is mainly expressed by epithelial cells of the lung, intestine, kidney, and colon (3). This may explain the high incidence of pneumonia and bronchitis in those with severe COVID-19 infection^5,14^. Analyses of the publicly available RNA data sets and immunohistochemical studies suggest that the ACE2 is also highly expressed in the skin and mucous membranes including the oral mucosa^15-17^. Consistently, vesiculobullous lesions affecting the skin and the oral mucosa have been reported as clinical presentations in COVID-19^18,19^. Since the viral attachment and entry is critical for replication and infection, the state of cell differentiation and the surface expression of ACE2 can directly influence the SARS-CoV disease pathogenesis. Hence, the goal of this study is to investigate the localization of ACE2 in mucosa from different regions of the oral cavity.

## Materials and methods

### Tissues

Archived paraffin embedded tissue blocks of buccal mucosa and tongue tissues with histological diagnosis of benign mucosal lesions such as fibroma and papilloma were obtained from the repository of the Oral Pathology Biopsy services at the Indiana University School of Dentistry. Hematoxylin and eosin stained sections were evaluated for the presence of sufficient amount of normal oral epithelium adjacent to the lesion in each tissue. Only tissues that possessed significant normal oral epithelium with no evidence of pathology were included for assessment of ACE2 expression.

Isolation of epithelial cells in saliva: Unstimulated whole saliva was collected by the drooling method for 5-10 minutes into a 15ml chilled centrifuge tube after obtaining informed consent as described^20,21^. All samples were centrifuged at 250g for 10 minutes at 4°C. The cellular sediment obtained was reconstituted in isotonic saline supplemented with two drops of Zap-O globin to lyse blood corpuscles and centrifuged at 1400rpm for 10min at 4°C. After washing in saline, the cell suspension was passed through a 20μ filter. The epithelial cell-enriched preparation was then assessed by light microscope for morphology. A thin smear of the cells was then maintained in a humid chamber and immunostained for ACE-2 along with the tissue section.

### Immunofluorescence

The paraffin-embedded sections were dewaxed and rehydrated by sequential incubation in decreasing alcohol concentrations. The sections were then subjected to heat induced antigen retrieval in Tris-EDTA buffer for 10 minutes. The slides were allowed to cool, washed with 1XTBS buffer and then incubated with 3% H_2_O_2_ for 10min to quench endogenous peroxidase activity followed by washing and incubation in 5% goat anti-serum to minimize non-specific binding. The sections were then incubated with the primary anti-human ACE2 mouse monoclonal antibody (1:500, Catalog number: 66699-1-Ig, Clone No.: 2F12A4, Proteintech, Chicago) overnight at 4°C^7^. After washing, the sections were incubated in dark with Alexa Fluor 488 conjugated goat anti-mouse secondary antibody (Jackson ImmunoResearch Inc., PA) at room temperature for 2h. Nuclei were stained with propidium iodide (1:1000, BD Biosciences, CA). The tissues were washed and mounted with ProLong™gold antifade mounting medium (Thermofisher, CA). Immunostained sections were scanned with a NIKON multiphoton microscope attached with a DS Ri2 Camera (NIKON Instruments Ind, Melville, USA).

## Results and discussion

Membrane bound ACE2 is the port of entry for SARS-CoVs to the target cells. ACE2 transcript and protein expression has been reported in several human tissues by reverse transcriptase polymerase chain reaction and immunohistochemistry respectively^5,14,16^. We observed strong ACE2 expression in the epithelial cells of the oral mucosa. The expression was higher in the keratinized surface epithelial cells than in the spinous or basal cells. This was more evident in the tongue tissues than that of the buccal mucosa (Fig 1A, D). Merged images with the nuclei staining propidium iodide (Fig 1B, E) showed prominent expression of ACE2 localized to the cell membranes (Fig 1C, F). The exfoliated epithelial cells in the saliva also exhibited significant ACE2 expression (Fig 2A). Merged images with nuclei staining propidium iodide (Fig 2B) shows that ACE-2 expression is observed on the membranes of the epithelial cells in saliva (Fig 2C).

**Figure 1:**
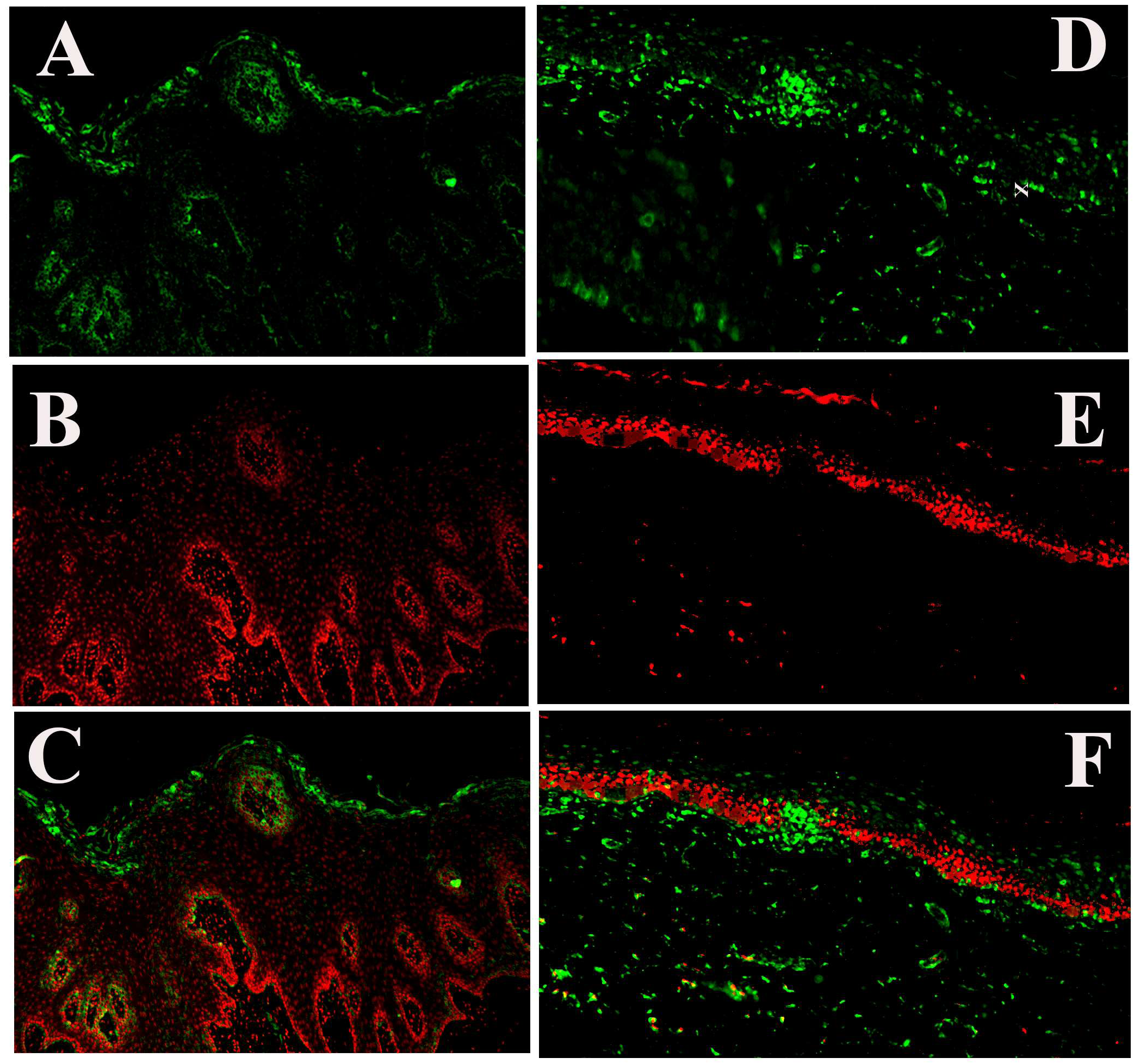
ACE2 is highly expressed in surface epithelial cells of the oral mucosa: Tongue and buccal mucosa biopsy tissues with adequate representation of normal oral epithelium were processed for immunofluorescence staining with anti-ACE-2 antibody (green) (A, C) and propidium iodide (red) (B, E) staining for nuclei. Positive staining for ACE2 is observed predominantly in the surface epithelial cells of the tongue (A, C) and buccal mucosa (D, F). Merged images (C, F) show that the ACE2 staining is seen in the cytoplasm and cell membrane.

**Figure 2:**
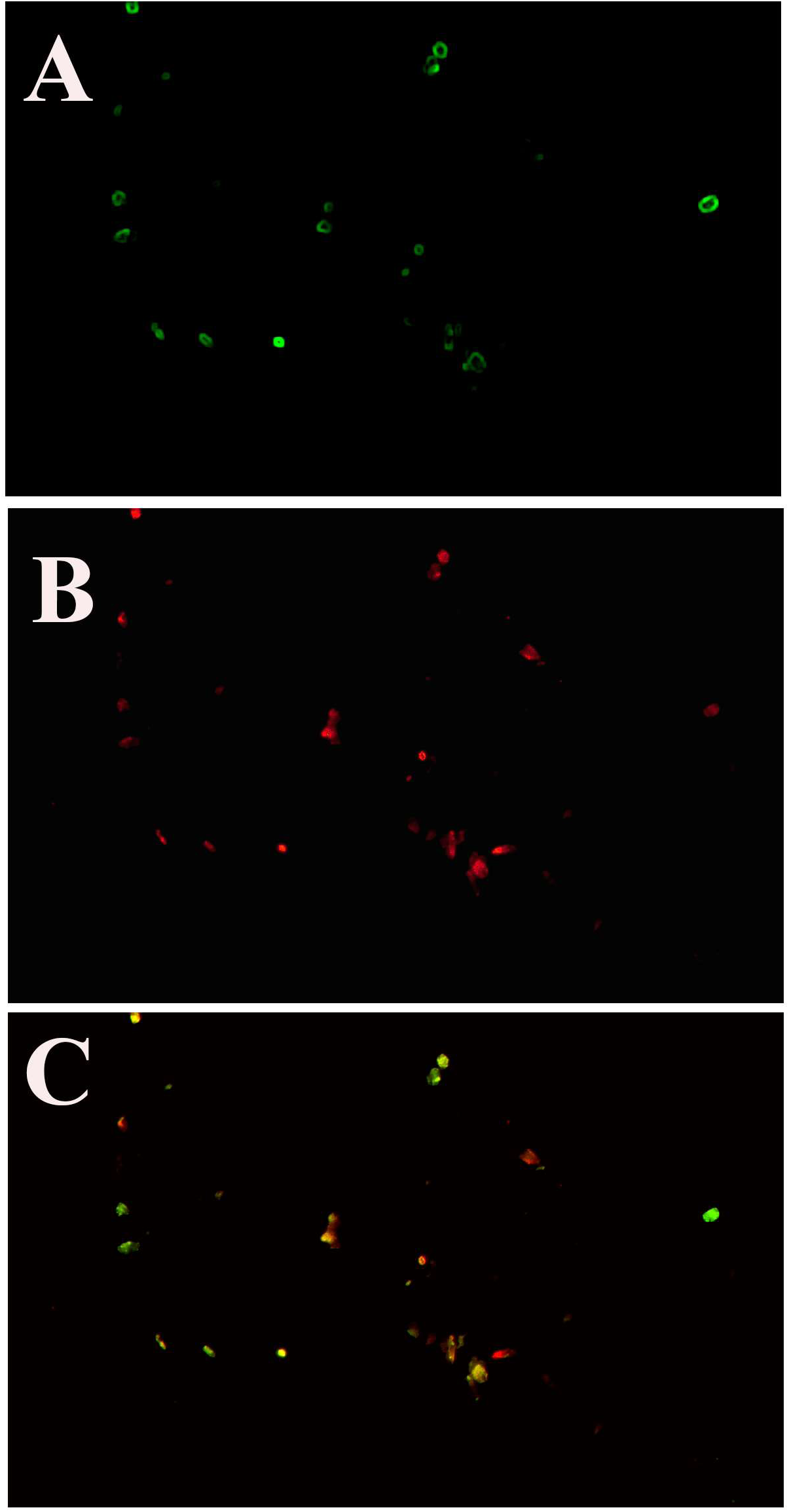
ACE2 is highly expressed in exfoliated epithelial in saliva. Oral epithelial cells isolated form unstimulated whole saliva were immunostained using fluorophore conjugated anti-ACE2 antibody (A) and propidium iodide (B) images at l0x and (C) shows merged image.

Epithelia are the primary barrier to microbial infection transmitted by contact with the exposed environment. As such, the epithelial cells express cell surface receptors that allow attachment to and penetration of the cell membrane by the microbes^20,22^. The distribution and density of the attachment receptors have been shown to be related with the degree of differentiation of the epithelia. Pertinent to SARS-CoV, the differentiated airway epithelial cells that abundantly express ACE2 have been shown to be more easily infected with the virus^7^. Indeed, in-vitro studies showed that SARS-CoV-1 virus enters by binding the ACE2 receptor abundantly expressed on the apical surface of the well-differentiated airway epithelia, especially the ciliated cells. Following replication, the virions are released from the basal surface of the infected cells^7,23^. In the skin, high expression of ACE2 has been observed in keratinocytes suggesting a percutaneous transmission potential for SARS-CoV^24^. Our observation of predominant expression of ACE2 in the surface epithelial cells of the tongue and buccal mucosa is consistent with these reports and support potential infection of the oral epithelial cells with SARS-CoV.

Structurally, ACE2 is a type I transmembrane protein with a short C-terminal cytoplasmic tail, a hydrophobic transmembrane region, and a glycosylated amino terminal ectodomain containing the active site^4,5^. A soluble form of ACE2 (sACE2) has been shown to be released either constitutively or induced by inflammation. The sACE2 arises from proteolytic cleavage of the ectodomain of the membrane bound ACE2 by the metalloproteinase ADAM-17, also known as TNF-α converting enzyme (TACE)^25^. Circulating levels of sACE2 has been shown to be low in children of both sexes, but increases with age, more so in healthy males than in females, so that as adults serum sACE2 is higher in men than in women ^26^.

In-vitro studies and animal experiments have shown critical roles for both membrane-bound ACE2 and sACE2 in the acute lung injury mediated by SARS-CoV^27^. The SARS-CoV spike protein not only binds the cell surface ACE2 but also has been shown to induce cleavage of ACE2 ectodomain together with the host protease TACE to achieve target cell entry^25^. The ACE2 shedding induced by the spike protein is tightly associated with the TNF-α production. Silencing of TACE with siRNA prevented viral entry suggesting that the SARS-spike protein induced TNF-α secretion and consequent TACE mediated ACE2 shedding is critical for infection^28^. Indeed, sACE2 has been suggested as a prognostic marker for SARS-CoV2 infection^29^.

In this context, our observation of ACE2 in the epithelial cells in saliva can be extended hypothetically as a potential marker for SARS-CoV-2 infection. It is likely that the surface epithelial cells of the oral mucosa that exhibit high expression of ACE2 act as initial sites of SARS-CoV-2 entry. Either the TNF-α induced by the viral attachment and/or released by the mild state of inflammation of the oral mucosa in health, may facilitate the SARS-spike protein mediated shedding of ACE2. Longitudinal observations in few SARS-CoV-2 positive cases have shown that the viral load in saliva was consistently higher during the symptomatic phase^30-32^. It is likely that the persistent infection could increase ACE2 ectodomain shedding from oral epithelial cells. Collectively, it can be postulated that the ACE2 levels in saliva could correlate with the SARS-CoV-2 viral load and potentially be predictive of symptom development in COVID-19 (Fig 3).

**Figure 3:**
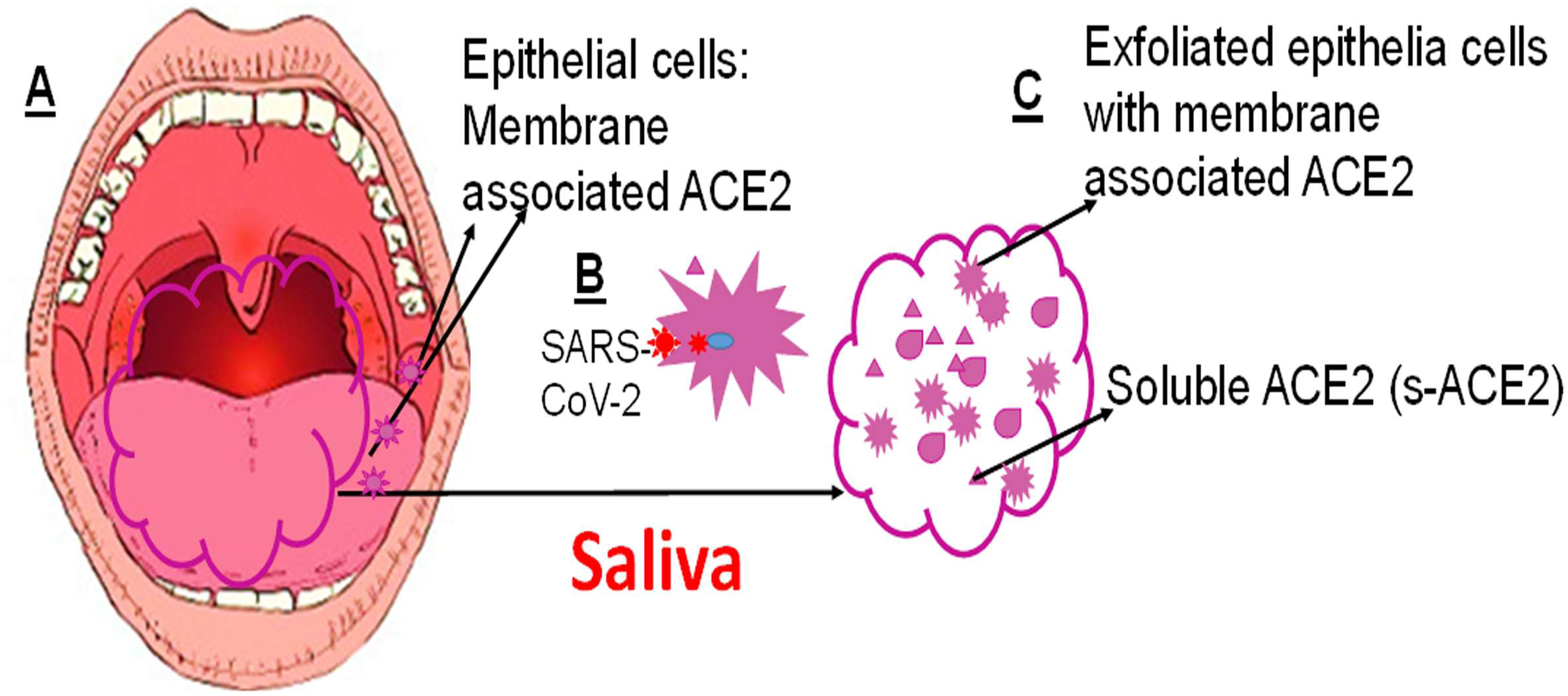
Schematic representation of potential value of assessment of ACE2 in saliva as a marker for SARS-Co V-2 infection and prognosis. ACE2 is the host receptor for SARS-Co V2. (A) The epithelial cells lining the oral mucosa express cell surface ACE-2, more so in the surface epithelial cells. (B) Upon binding the ACE2 on host cell surface, the spike protein of the SARS-Co V-2 activates the host proteases such as TACE that cleaves both the spike protein and the ACE2 at the transmembrane region. The virus then enters the cell and the ACE2 ectodomain is released as soluble ACE2 (sACE2). (C) The saliva is a rich source of exfoliated oral epithelial cells. In addition, saliva proteome includes as many as 5000 proteins that include secretions of the salivary gland, factors released by the epithelial cells, proteins reaching the saliva by passive diffusion from blood and microbial proteins. Taken together the membrane associated ACE2 in the exfoliated epithelial cells and the sACE-2 arising from ectodomain shedding locally or by passive diffusion from blood in saliva would be elevated in active SARS-Co V-2 infection. Assessment of ACE2 transcript together with the measurement of viral load in saliva represents a potential strategy to predict active infection and/or transmissibility of SARS-Co V-2.

## Conclusions

In a review of over 700 publications on diagnostic approaches of viral infections, the authors observed that the application of non-invasive sampling methods such as the use of saliva for diagnosis and monitoring of viral diseases need to be widely investigated^33^. This is particularly critical for respiratory infections such as SARS-CoV-2, in which the viral load in saliva has been shown to be higher in early phase of the disease^31,32,34^. Based on our histological observations and the rationale discussed, we postulate that the assessment of SARS-CoV-2 viral load and ACE2 in the saliva could represent potential biomarkers for COVID-19 diagnosis and prognosis.

## Competing interest

The authors declare no competing interest.

## Notes

### Competing Interest Statement

The authors have declared no competing interest.

